# Disentanglement of persistent and dynamic neural substrates of working memory

**DOI:** 10.64898/2026.11.05.764743

**Authors:** Alireza Abbasi, Aditya Nanda, Wenhao Dang, Balbir Singh, Zhengyang Wang, Christos Constantinidis, Mika Rubinov

## Abstract

Competing theories of working memory have alternatively emphasized the importance of persistent and dynamic substrates of this cognitive function. Here, we directly tested these theories by computationally isolating the contributions of persistent and dynamic activity to working memory. We analyzed neuronal population activity recordings in macaques doing match/non-match working memory tasks. We first extracted signatures of persistent and dynamic activity through cue-specific temporal means, temporal principal components, and correlations of activity patterns across task epochs. We then generated synthetic data constrained by these signatures, and decoded cue stimuli from these data, separately from persistent and non-persistent neurons. We found that high decoding performance arose from a nuanced interplay of persistent and dynamic activity. Specifically, we could decode stimuli at empirical or above-chance levels across all tasks only from persistent neurons, but only after constraining correlations of activity patterns across epochs. In this way, we showed distinct roles for persistent and dynamic neuronal activity that help to unify two competing theories of working memory. More broadly, we also advanced computational controlled experiments as a principled frame work for adjudicating between competing models of neuronal population activity.

## Introduction

Working memory helps humans and animals maintain internal activity patterns over brief periods (1). The neural substrate of this memory remains an open question. Important early studies have linked this substrate to patterns of neuronal activity in the prefrontal cortex (2). These studies have specifically shown that some neurons in the prefrontal cortex are active during delay periods, after the removal of cue stimuli (3–5). In what follows, we use the term *persistent activity* to describe these patterns of activity and *persistent neurons* to describe the neurons that show these patterns of activity.

The relationship between persistent activity and working memory has been an object of longstanding interest, but also some debate (6,7). On the one hand, studies have shown the presence of persistent activity across species and experiments (8,9). In parallel, modeling work has shown that stimulus specific activation of recurrent network models can generate persistent activity as a continuous attractor state (10,11). On the other hand, studies have also successfully decoded cue stimuli from *non-persistent* neurons that do not have elevated firing rates during delay periods (12). On this basis, alternative theories have proposed that networks can keep short-term activity through non-persistent or dynamic mechanisms, including synaptic plasticity (13–15). Studies have sometimes termed these latter mechanisms as “activity-silent”.

We recently compared these theories by showing that we could decode cue stimuli much better from persistent rather than non-persistent neurons (16). Such comparisons, however, ultimately remain observational and make it difficult to disentangle the contribution of distinct types of neurons and activity to working memory.

Here, we adopted a computational framework that goes beyond observational evidence to directly test the effects of persistent and dynamic activity on decoding performance (17–19). The framework broadly parallels a randomized controlled trial (19) and consists of three steps. In the first step, we defined statistics of the data that captured signatures of persistent or dynamic neuronal population activity. In the second step, we generated samples of synthetic data that satisfied these statistics but were maximally random in all other respects. In the third step, we tested the decoding performance of persistent and non-persistent neurons in the individual data samples. This approach thus allowed us to test, in a controlled way, whether statistical signatures of persistent or dynamic population activity were sufficient to decode cue stimuli, separately for persistent and non-persistent neurons.

We applied this approach to neurophysiological recordings from the prefrontal cortex of non-human primates performing three types of working-memory tasks. We built models that encode aspects of persistent activity (as cue-specific means or principal components), and dynamic activity (as pairwise correlations of mean-activity patterns across task epochs). We also used weaker control models to more precisely disentangle the nature of the main effects. Together, this controlled approach allowed us to simultaneously pinpoint neurons that helped keep cue structure, and pinpoint aspects of neuronal activity that helped them do so across cognitive tasks.

## Results

We analyzed neuronal activity from the lateral prefrontal cortex of three rhesus monkeys trained to do match/non-match working memory tasks. In each trial, the monkeys first viewed two stimuli (cue and test) separated by a delay period and then had to decide, after 1.5 seconds, if the test stimulus matched the cue stimulus. The monkeys signaled their decision by making a saccade to one of two color-coded targets. In case of matching cue and test stimuli, monkeys were trained to saccade to a green target; otherwise, they were trained to saccade to a blue target located at a diametrically opposite position. We studied three tasks with distinct match conditions: in a spatial task, the stimuli matched in location; in a feature task, they matched by shape; and in a conjunction task, they matched by both location and shape. The schematics in **Figure S1** show the structure of these tasks.

We analyzed the activity of 1,062 neurons across the three animals for the spatial task, 884 neurons for the feature task, and 484 neurons for the conjunction task. For downstream analyses, we normalized the activity of each neuron to have zero mean and unit variance. To avoid the possibility of “double dipping” (20) or “data leakage” (21), we did this normalization separately within each trial.

We then classified the neurons and time points in the following way:

- We classified all neurons into two activity-dependent groups. We defined neurons to be *persistent* if they showed significantly elevated firing rates during the first delay period. Correspondingly, we defined the neurons to be *non-persistent* if they did not show elevated firing rates during the first delay period — either because they showed no elevation in firing rate during any epoch, showed lower firing rates during the first delay period, or showed elevated firing only during epochs other than the first delay period. The number of persistent neurons was 413 (38.88%) in the spatial task, 315 (35.63%) in the feature task, and 273 (56.40%) in the conjunction task.
- We labeled every time point in each recording in several ways:

∘ By the nature of the task (spatial, feature, conjunction).
∘ By the specific cue stimulus within each task (eight locations for the spatial task, eight shapes for the feature task, four combinations of locations and shapes for the conjunction task).
∘ By the trial number for each task (1–12 trials for spatial and feature tasks, 1–24 trials for the conjunction task).
∘ By the task epoch within each trial (pre-fixation, fixation, cue stimulus, first delay, test stimulus, second delay, response).

We used this classification to model the activity of distinct neurons across tasks, stimuli, trials, and epochs in a consistent and interpretable way. **Figure 1** shows the structure of this classification.

**Figure 1.**
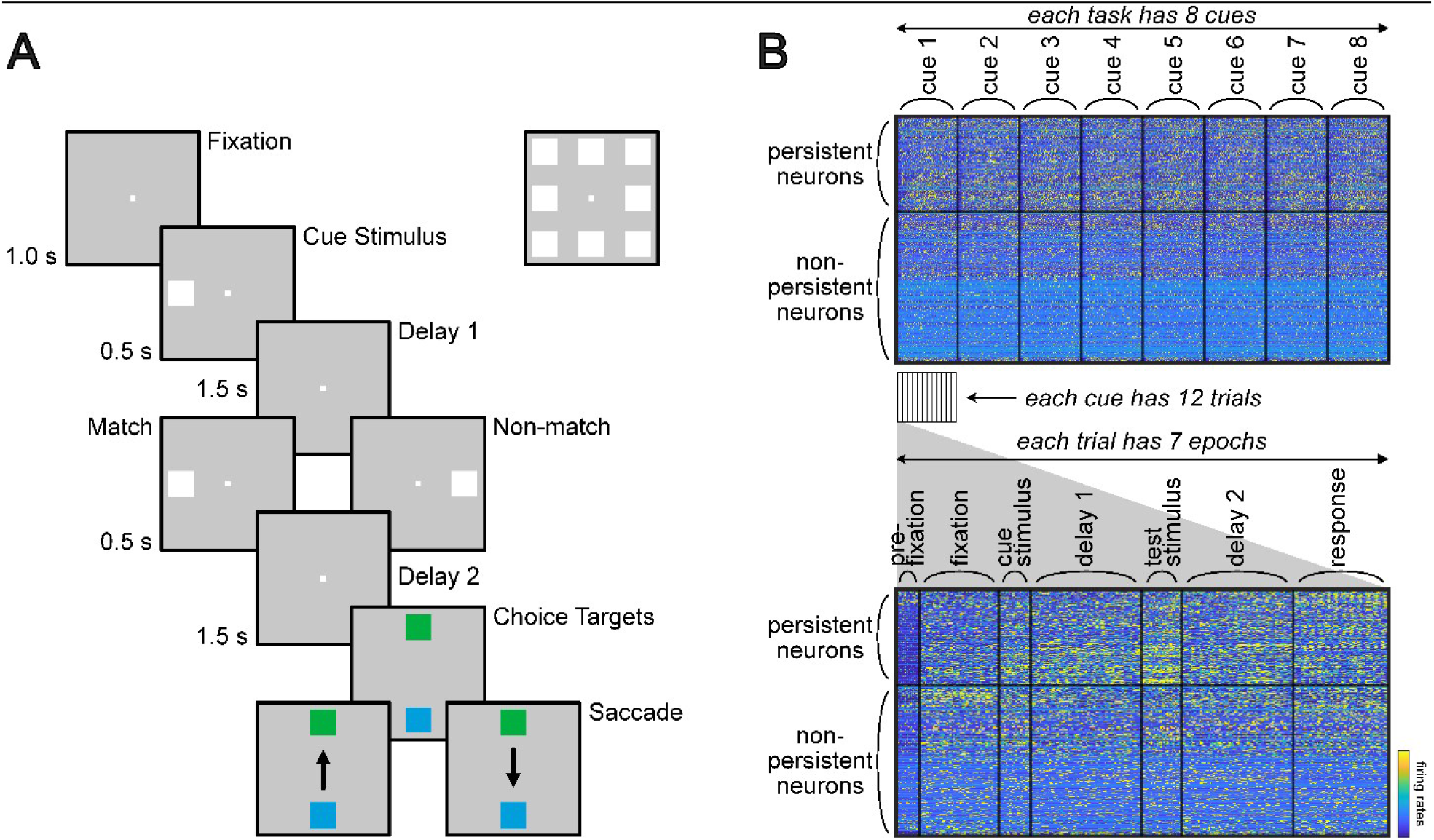
Structure of task and neuronal population activity. **A**. Structure of the spatial task. **B**. Schematic of neuronal population activity matrix for each task. Rows in this matrix rep resent neurons, while columns represent time points. The neurons are split into persistent and non-persistent categories. The time points are split into either 8 cues × 12 trials (for spatial and feature tasks) or 4 cues × 24 trials (conjunction task). Each trial has seven task epochs: pre-fixation, fixation, cue stimulus, first delay, test stimulus, second delay, and response. Colors represent firing rates at individual time points.

### A framework for statistical modeling of working memory

We began our analyses by defining statistical models of persistent and dynamic activity across tasks, trials, and epochs. All our models were based on statistical signatures that we estimated directly from the data, separately for persistent and non-persistent neurons.

1. We defined models that preserved the low-dimensional structure of population activity across time. These models captured delay-related increases in firing characteristic of persistent activity. We considered two variants of these models: *Tuned-cluster mean model (cluster model).* This model preserved the mean neuronal activity within clusters of neurons, with each cluster containing neurons most sensitive or tuned to individual cues. This model, therefore, captured delay-specific elevation in activity associated with persistent firing within tuned neuronal groups **(Figure 2B)**. *Temporal component model (component model).* This model preserved the mean neuronal activity and the first three temporal principal components for each trial. This model, therefore, captured the dynamic temporal structure in population activity, beyond the overall delay-related increase in firing characteristic of persistent neurons **(Figure 2A)**.
2. Separately, we defined models that preserved pairwise correlations between mean activity patterns within epochs, or whole trials. These models thus captured coordinated relationships between neuronal population activity but did not necessarily preserve mean activity or other low-dimensional features characteristic of persistent activity. We considered three variants of these models: *All-epoch correlation model (all-epoch model).* This model preserved correlated activity between all pairs of epochs across all stimuli and trials. For spatial and feature tasks, the dataset comprised 8 cues with 12 trials per cue, while for the conjunction task it comprised 4 cues and 24 trials per cue **(Figure 1)**. Each trial comprised 7 epochs, which for all tasks resulted in a 672 x 672 correlation matrix **(Figure 2C)**. *Same-epoch correlation model (same-epoch model).* This model captured correlated activity be tween all pairs of epochs of the same type. For each task, this model thus effectively preserved seven 96 × 96 correlation matrices, i.e., one correlation matrix for each epoch **(Figure 2C)**. *Whole-trial correlation model (whole-trial model).* This control model preserved the correlation structure of trial-averaged activity across all seven epochs. For each task, this model thus effectively preserved one 96 × 96 correlation matrix of all trials **(Figure 2D)**.

**Figure 2.**
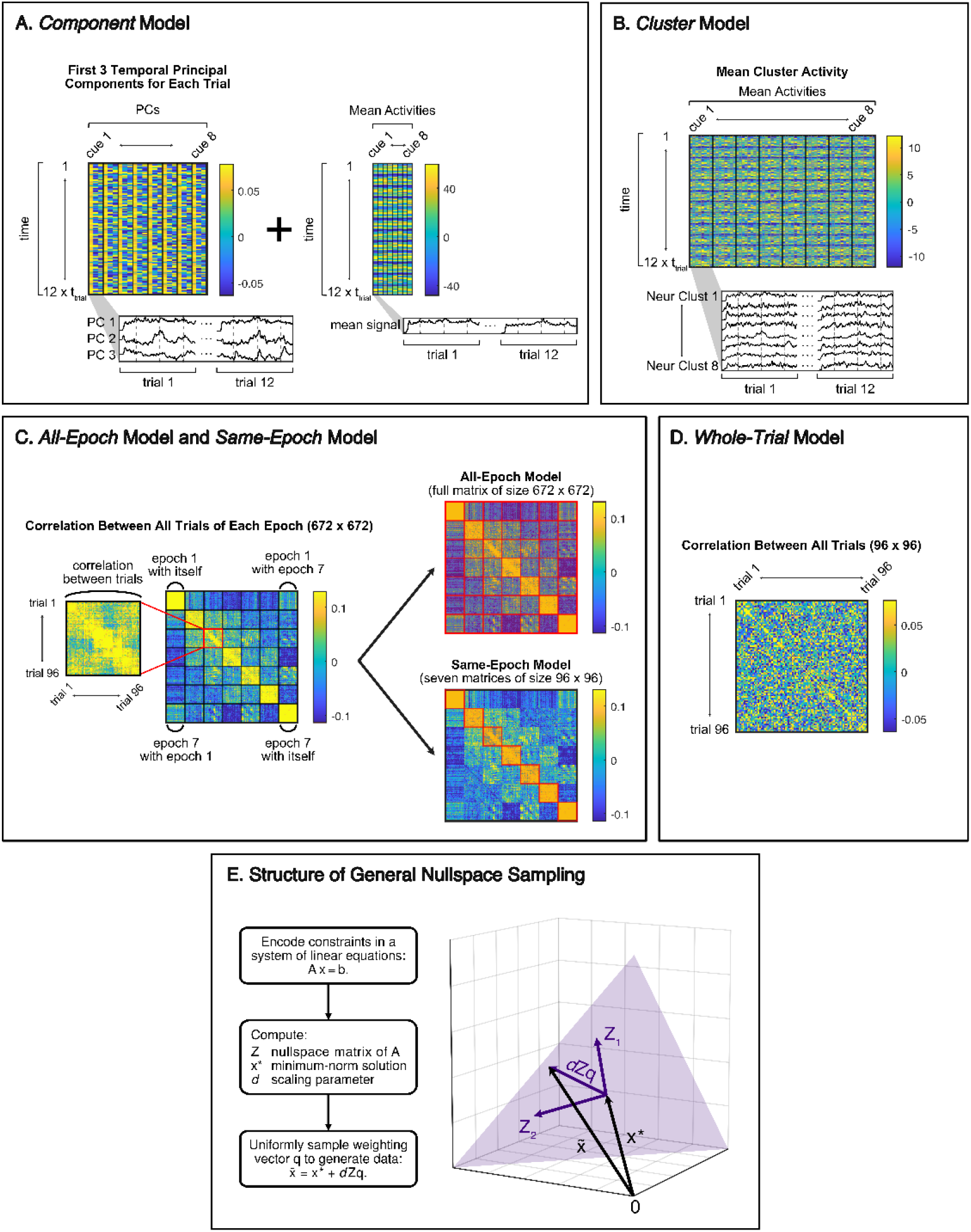
Model definition and general structure of nullspace sampling. **A**.A matrix of the first three principal components (left) and mean activities (right) across stimuli and trials. Each column represents a principal component (left) or mean activity (right), and each column corresponds to the 12 trials of each cue. The component model preserved these components and mean activities separately for persistent and non-persistent neurons. **B**.A matrix of the mean activities of each preferred-cue neuronal cluster, across stimuli and trials. Neurons were assigned to one of the clusters by the cue eliciting their largest cue-period response relative to fixation, so each column of panel A expands into eight cluster means (four means for the conjunction task). The cluster model preserved these activities separately for persistent and non-persistent neurons. **C.Left**. Matrices of correlations between activity patterns across pairs of 672 epochs (7 epochs/trial × 12 trials/cue × 8 cues for the spatial and feature tasks, or 7 epochs/trial × 24 trials/cue × 4 cues for the conjunction task). The matrices are first ordered by epoch type, from pre-fixation to response (the 7 diagonal blocks), and then by the order of trials (inset). **Right**. The all-epoch model preserved correlations between all pairs of epochs (top), while the same-epoch model preserved correlations between epochs of the same type (bottom). Both models preserved correlations separately for persistent and non persistent neurons. **D**.Matrix of correlations between trial-averaged activity patterns across pairs of 96 trials. The whole-trial model preserved these correlations separately for persistent and non-persistent neurons. **E**.General formulation of nullspace sampling method. The algorithm first encodes statistical constraints in systems of linear equations. It then sequentially generates synthetic data using the nullspace of this system. See Methods for additional details.

After defining each model in terms of the above statistics, we generated synthetic data constrained by these statistics and estimated cue decoding in these model data against empirical decoding. In this way, our approach essentially formed a computational controlled experiment for testing the decoding performance of individual statistical signatures, and by extension, the associated persistent or dynamic aspects of neuronal activity during working-memory tasks.

We generated synthetic data using nullspace sampling, a method that we recently developed for testing competing explanations of brain activity data (22,23). The method allowed us to translate a wide range of neurobiological signatures, including temporal means, principal components, and pattern correlations, into sets of systems of linear equations. The method uses principles of linear algebra to sample the space of all solutions that satisfy these equations and thus generate data constrained by these properties but maximally random in all other respects. Figure 2E shows a schematic of this method, while the Methods section describes it in more detail.

### Statistical signatures and decoding performance

**Figure 3** shows the pertinent statistical features of empirical data and the corresponding model data defined in the previous section. (see **Figures S2–S8** for illustration of all features). It specifically shows the temporal mean **(Figure 3A)** and principal components **(Figure 3B)** and, separately, the all-pair wise correlations of activity patterns across task epochs **(Figure 3C)**. The figure shows that no individual model preserved all these features simultaneously. Specifically, the cluster and component models lacked the rich structure of dynamical correlations, while the correlation models lacked the empirical mean and principal components of the individual trials. Together, the full compendium of these models thus allowed us to do targeted tests of the importance of individual constraints on the decoding of cue stimuli.

**Figure 3.**
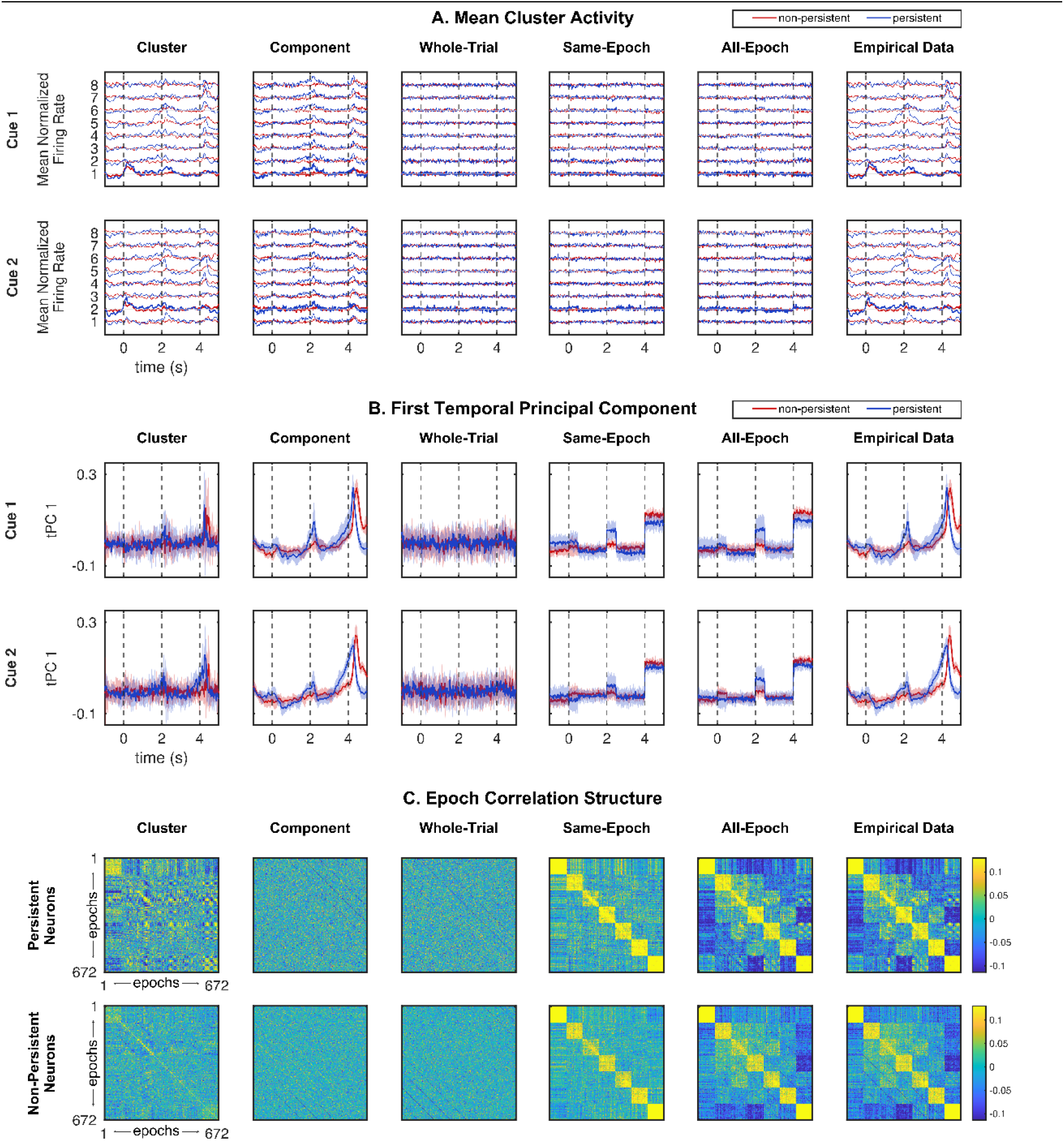
Statistical features of competing models and empirical data for the spatial task. Columns 1–2 show the cluster and component models. Columns 3–5 show the three correlation models. Column 6 shows empirical data. **A**.Mean normalized firing rate for each of the eight preferred-cue neuronal clusters, shown for persistent and non-persistent neurons for two representative cues. Traces are offset vertically by cluster, and the cluster matching the cue of that row is drawn in bold. **B**.First temporal principal component, shown for persistent and non-persistent neurons for two representative cue stimuli, aligned to a common reference by Procrustes analysis. The shaded area shows the standard deviation of these values across all 12 trials. Dashed lines at 0, 2, and 4 s denote the onsets of the cue, the test stimulus, and the response epoch, respectively. **C**. Pairwise correlations between mean activities within each epoch across all trials in the spatial task, shown separately for persistent and non-persistent neurons and ordered as in **Figure 2C. Figures S2–8** show a comprehensive illustration of these constraints across all tasks and cues.

We next evaluated how well the activity of persistent and non-persistent neurons could decode cue stimuli. We did this by randomly selecting 200 neurons from either the persistent or non-persistent populations, training a support vector machine on the activity of these neurons at specific time intervals to classify stimulus cue identity, testing this performance with 10-fold cross-validation, and repeating this process 100 times.

**Figures 4 and 5** show our main results for the model and empirical data across the three tasks. **Figure 4** shows the decoding performance at specific time points, based on a sliding window of 100 ms with a step size of 25 ms. Complementarily, **Figure 5** shows decoding performance during the middle 1 second of the delay period, and thus more directly captures the relevance of these results to working memory. The delay period spans 0.5 to 2 seconds after cue onset, and we used the decoding window from 0.75 to 1.75 seconds. This choice minimized potential carryover effects from cue presentation and ensured that the decoding reflected delay-period activity specifically. We quantified this performance as medians [90% confidence intervals], and considered a result to be significant at *p* < 0.05 if the lower bound of this interval exceeds the chance level, which was 0.125 (1/8 stimuli) for the spatial and feature tasks and 0.25 (1/4 stimuli) for the conjunction task. In addition, Figure 5 also shows the relative number of constraints for each model. Models that have fewer constraints correspond to simpler representations of neural population structure and, all else being equal, are preferred by the principle of parsimony.

**Figure 4.**
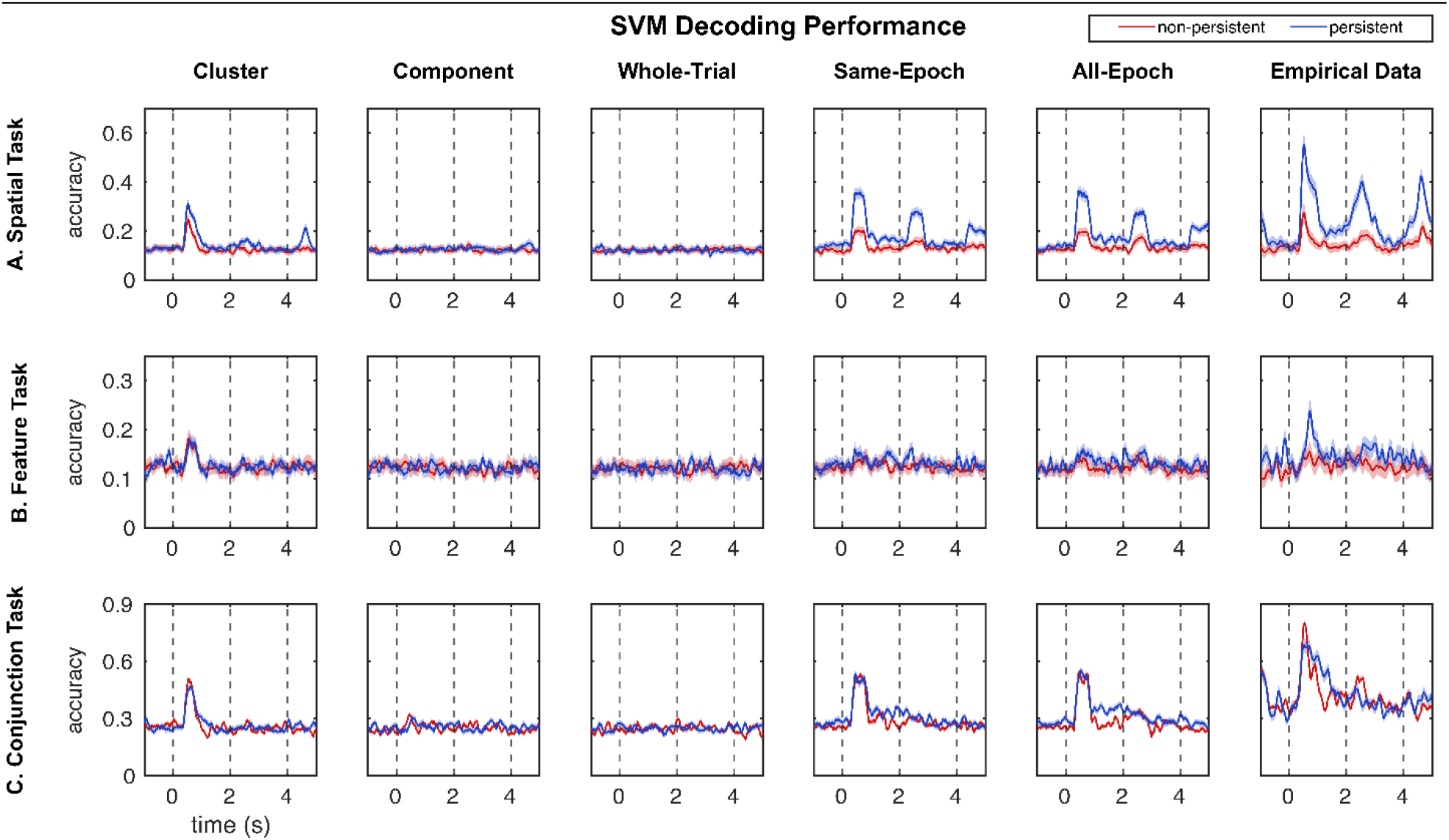
Cue decoding performance for empirical and model data. Columns 1–2 show the cluster and component models. Columns 3–5 show the correlation models. Column 6 shows empirical data. Dashed lines at 0, 2, and 4 s denote the onsets of the cue, the test stimulus, and the response epoch, respectively. **A**.SVM decoding performance for the spatial task as a function of time, calculated using a 100-ms window and a 25-ms step size. Cue decoding accuracy was computed using 200 neurons randomly sampled from each neuronal category (persistent or non-persistent) with 10-fold cross-validation repeated over 100 iterations. The shaded area indicates the standard deviation of accuracies across iterations, and the curves represent the mean decoding accuracy over all 100 iterations. Accuracies were calculated separately for persistent and non-persistent neurons. **B**.SVM decoding performance for the feature task, calculated using the same process as in panel **A. C**.SVM decoding performance for the conjunction task, calculated using the same process as in panel **A**.

**Figure 5.**
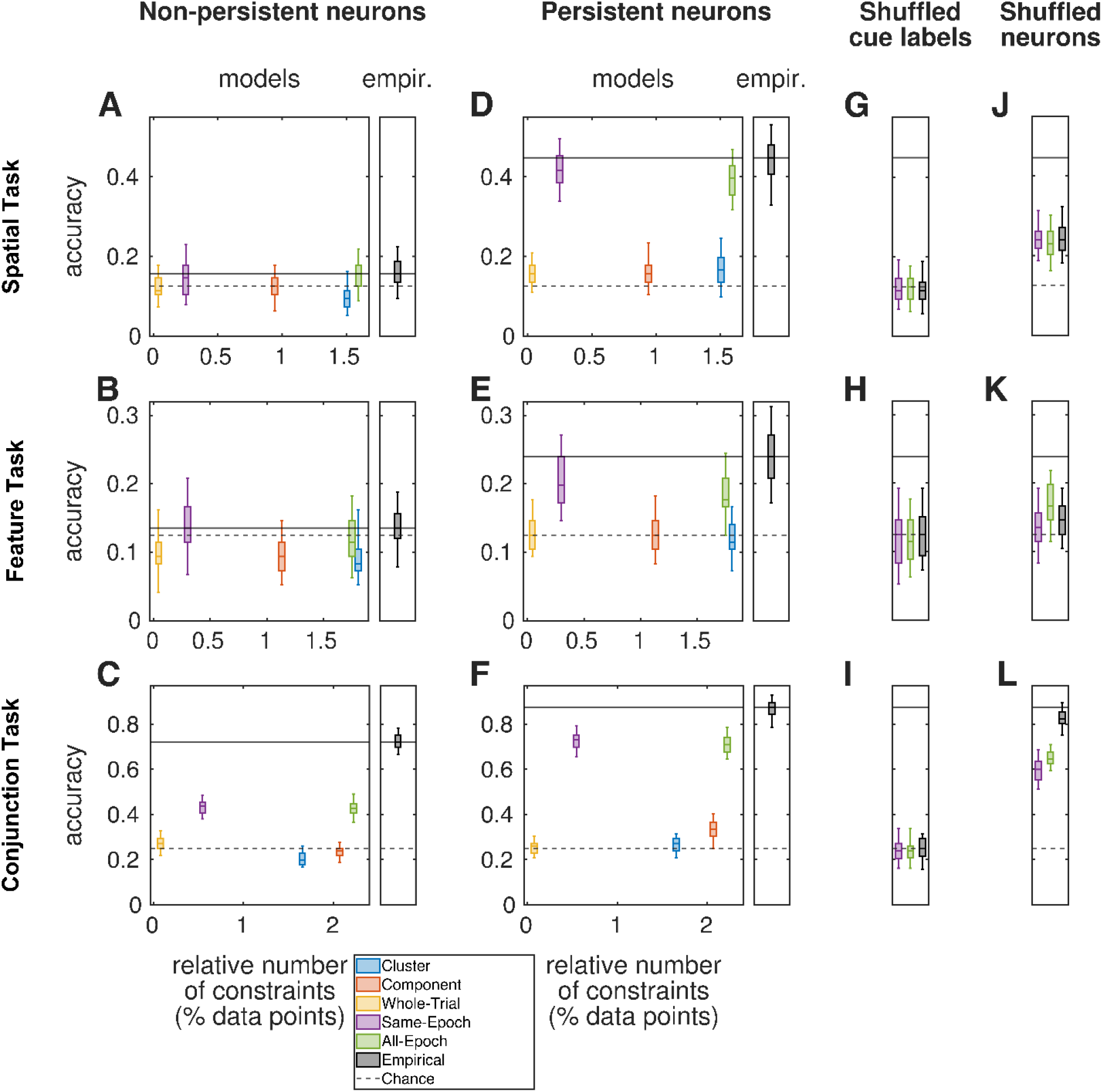
Cue decoding performance during the delay period for models and empirical data. Box plots show cue SVM decoding performance computed using 10-fold cross-validation over 200 randomly selected neurons for 100 iterations. Models were trained using the middle 1-second window of the delay period (from 0.75 to 1.75 seconds after cue onset) during all three tasks. Note that these values are not equivalent to a simple average of decoding performance across individual time points shown in **Figure 4**. Boxes denote the first and third quartiles, and the whiskers denote the 90% confidence interval. In panels A-F, the horizontal axis shows the total number of constraints as percentages of the empirical data. The dashed line represents chance accuracy (12.5% for the spatial and feature tasks, 25% for the conjunction task), and the solid line represents the median accuracy for the empirical data. **A–C**. Decoding from non-persistent neurons in the model and empirical data across the three tasks. **D–F**. Decoding from persistent neurons in the model and empirical data across the three tasks. **G–1**. Decoding from persistent neurons in the correlation models and empirical data after shuffling cue labels across the three tasks. **J–L**. Decoding across the three tasks from the correlation models and empirical data after shuffling neuron labels across the three tasks.

We previously found that empirical cue decoding accuracy during the delay period of the spatial and feature tasks was high for persistent neurons (16). **Figure 4 and 5** reproduced these results across the three tasks. In this way, we found that the decoding level of persistent neurons in the empirical data was above chance level for all tasks: 0.448 [0.328, 0.531] for the spatial task, 0.234 [0.172, 0.313] for the feature task, and 0.875 [0.787, 0.927] for the conjunction task. By contrast, non-persistent neurons performed substantially worse in all three tasks, with values of 0.156 [0.094, 0.224] for the spatial task, 0.135 [0.078, 0.188] for the feature task, and 0.719 [0.667, 0.781] for the conjunction task—at chance level for the spatial and feature tasks, and significantly below persistent neurons in the conjunction task.

One superficial interpretation of these results is that high decoding performance arises simply from the mean activity differences or other low-dimensional features of persistent neurons. This interpretation, while intuitive, was not consistent with our modeling results. Specifically, we found that the cluster and component models accurately captured the structure of empirical persistent activity **(Figure 3A-B)**, and yet did not exceed chance-level decoding. For example, persistent neurons from the component model had decoding performances of 0.156 [0.104, 0.234], 0.125 [0.083, 0.182], and 0.333 [0.250, 0.401] for the three respective tasks (Figure 5). These results imply that the structure of cue stimuli was insufficiently encoded in the clusters or components of the population activity in our data.

By contrast, persistent neurons in the two models that preserved correlations between all epochs (all epoch model), or only epochs of the same type (same-epoch model), showed above-chance decoding performance, even if the synthetic timeseries of these neurons lacked the outward manifestations of persistent activity (in other words, these neurons are persistent in the empirical data, but not in the synthetic data, **Figure 3A)**. Specifically, we found that the decoding performance for persistent neurons in the all-epoch model was 0.396 [0.318, 0.469] for the spatial task, 0.177 [0.125, 0.245] for the feature task, and 0.708 [0.646, 0.786] for the conjunction task. Notably, the performance of the considerably simpler same-epoch model was similar at 0.417 [0.339, 0.495] for the spatial task and 0.198 [0.146, 0.271] for the feature task, and 0.729 [0.656, 0.792] for the conjunction task. Also notably, and as in the empirical data, the performance of non-persistent neurons in these model data was consistently lower than that of persistent neurons, and did not exceed chance in the spatial and feature tasks. Finally, the even simpler model, which collapsed activity across all epochs in the trial (whole-trial model), did not achieve above-chance decoding, with corresponding performance of 0.156 [0.109, 0.208], 0.125 [0.094, 0.177], and 0.260 [0.208, 0.302] for the three respective tasks.

We next sought to disentangle the drivers of this performance in three ways:

First, we randomly permuted the identity of individual cue stimuli, while preserving all other aspects of the data. This test helped us see whether the performance of the correlation models reflected empirical encoding of cues, rather than the idiosyncratic trial-specific structure. In the former case, we expected this permutation to lead to chance-level performance, while in the latter problematic case, the performance would not be affected and thus stay artificially high. In practice, we found that per-mutation led to degraded performance in both empirical and model datasets **(Figure 5G–1)**. This pro vided complementary evidence for the importance of these model constraints for decoding accuracy.

Second, we randomly permuted the persistent and non-persistent labels across neurons, while pre serving the size of each neuronal group. This test helped us see whether high decoding accuracy was specific to persistent neurons. In practice, we found that the permuted groups showed intermediate performance, most clearly in the spatial task **(Figure 5J–L)**. This confirmed that the correlation structure driving cue decoding was specific to the persistent-neuron population.

Finally, we directly tested if statistics of individual models can distinguish between cues in empirical data. To do this, we extracted epoch means, temporal components, and cluster means from empirical data, and estimated the difference in similarity between these statistics between- and within-cues (Figure 6). We found that the epoch mean statistic showed the largest cue-specificity, compared to cluster means and temporal components. These results align with our observed decoding performance and help to more directly establish the statistical underpinnings of this performance.

**Figure 6.**
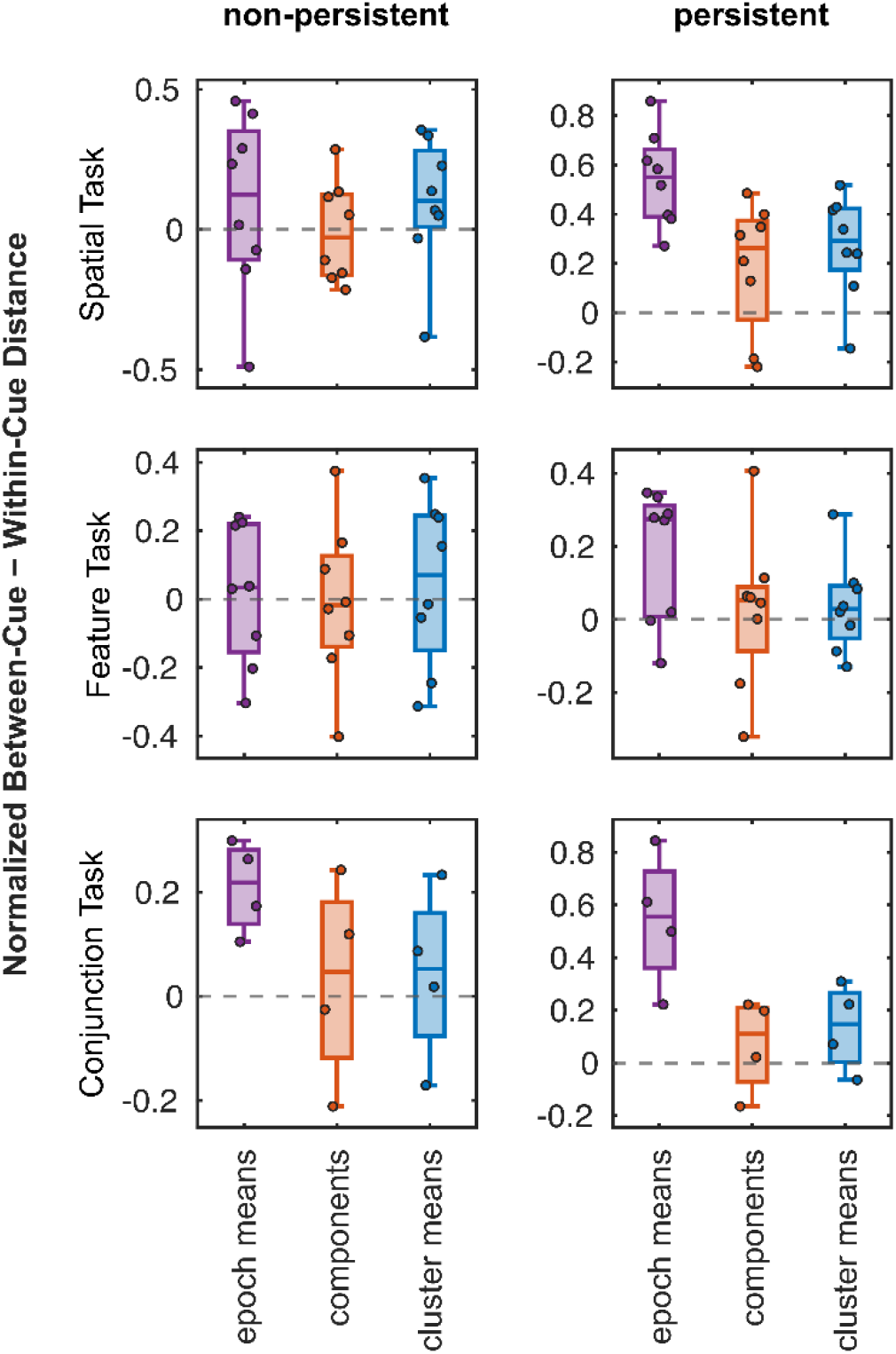
Between- and within-cue distances for individual model constraints. Box plots show differences between between-cue and within-cue trial distances in the empirical data during the middle 1-second window of the delay period (from 0.75 to 1.75 seconds after cue onset) for the three statistics preserved by the models: epoch means, principal components, and cluster means. Rows correspond to the spatial, feature, and conjunction tasks, and columns correspond to non-persistent and persistent neurons. For each statistic, we normalized pairwise squared Euclidean distances between trials and computed the difference between the mean between-cue and within-cue distances (positive values indicate smaller relative distances). Points represent individual cues (eight for the spatial and feature tasks, four for the conjunction task). Boxes denote the first and third quartiles, whiskers denote the 90% confidence interval, and the dashed line represents zero (equal between- and within-cue distances).

## Discussion

We adopted a principled statistical modeling framework to study the structure of population activity in the prefrontal cortex during working memory. We modeled recordings of spatial, feature, and con-junction working memory tasks by generating data that preserved persistent and dynamic features of neuronal activity, separately for persistent and non-persistent neurons. Our results show a nuanced interplay of aspects of neuronal activity. On the one hand, we showed that persistent neurons encode cue stimuli in empirical data and that this signal is not captured by cluster and component models. This suggests that the persistent population disproportionately contributes to the formation and maintenance of structured task representations. On the other hand, we found that the main statistical features that facilitated this formation and maintenance were the cue-specific patterns of correlation between mean activity within task epochs, rather than stable “persistent” increases in firing rate dur ing the delay period. Therefore, persistent neurons can help encode stimuli through dynamic effects, including rich correlation structures, which prefrontal neurons are known to exhibit (24,25).

The detection of activity during the delay period in the prefrontal cortex has generally been an uncontroversial hallmark of working-memory studies over the last thirty years (2) and recent experiments have provided critical validation of persistent activity across the population of prefrontal neurons, in single trials (26,27). Some studies have questioned whether this activity is persistent or whether it sporadically varies across trials, or across the delay period within individual trials (7). In our task, robust persistent activity was present for all aspects of information maintained in working memory (spatial location, shape, and their combination). However, several memory phenomena have been described in which persistent activity is absent or greatly diminished (28,29). For example, serial bias is a phenomenon in which the location of a previous trial’s stimulus influences the response in the current trial, even though persistent activity does not survive between successive trials. It is clear, there-fore, that non-spiking mechanisms, such as short-term synaptic modification or intracellular calcium availability, contribute to such mechanisms (28,30).

Our results help clarify this question by noting that a class of persistent neurons that show such activity in their firing rates keeps cue structure, but through distinct mechanisms that center on pattern specific correlations. Mathematically, “persistent” and “dynamic” constraints in our study roughly de-note distinct approximations of all-pairwise mean activity patterns across time points of the full re cording. Specifically, the cluster and component models represent the means or eigenvectors of the trial-specific portions of this matrix, while the correlation models roughly represent within-epoch or within-trial averages of portions of this matrix. In this way, the study ultimately blurs distinctions between persistent-activity and dynamic paradigms. Our observation that persistent neurons are the specific population carrying information, combined with the finding that they do so through epoch correlations, in effect proposes a hybrid model that supports complementary aspects of both mechanisms, where persistent subpopulations support working memory via cue-specific correlation pat terns, rather than through cue-specific firing levels alone. Our findings also open important questions about the interaction of persistent and non-persistent neuronal populations, the limits of dynamic mechanisms, and the generalization to other tasks or brain regions.

### Limitations and future directions

Our study is not without limitations. First, we did not attempt to distinguish the specific patterns of dynamic activity that give rise to the correlations we constrained in this study. In that respect, our approach does not differ from previous studies of activity-silent populations which have also focused on decoding performance, without seeking to extract mechanisms (15). Careful future experiments could, however, test these mechanisms by manipulating the activity patterns of these neurons. Second, our analysis relied on pseudo-populations of neurons recorded at different sessions but treated as synchronous. The real-time structure of neuronal firing during task execution is weakly correlated between neurons (31,32), imposing additional constraints on the correlation structure of the population. As recordings from increasing numbers of neurons in primates are becoming possible (27,33), this limitation, too, can be addressed in future studies.

Finally, we note that the study used nullspace sampling to generate model data with diverse persistent and dynamic constraints. We propose that advances in neurobiologically informed synthetic data generation methods will become increasingly important for modeling of these questions in neuroscience. Ultimately, we consider that widespread adoption of such methods can help facilitate unified explanations of brain activity across complex cognition and behavior (19).

## Methods

### Experimental Setup

We collected neural recordings from three male rhesus monkeys (Macaca mulatta) performing working memory tasks. The monkeys were aged 5–9 years, weighed 5–12 kg, and were housed individually or in pairs with sensory access to other monkeys. All procedures complied with institutional and national animal care guidelines (34).

During the recordings, the monkeys were head-fixed in a primate chair and viewed a screen 68 cm away in a dimly lit room. A central 0.2° fixation target was used to stabilize gaze, and visual stimuli were presented either centrally or peripherally. Monkeys were rewarded with fruit juice for maintaining fixation, and the trials were aborted if fixation was broken. Eye position was tracked using a high resolution infrared system sampled at 240 Hz.

Neural activity was recorded from both dorsal and ventral regions of the prefrontal cortex using arrays of 1–8 glass- or epoxylite-coated tungsten microelectrodes (200-250 μm diameter, 1–4 MΩ impedance) advanced into the cortex with a motorized microdrive. Signals were amplified, filtered, digitized, and spike-sorted offline using principal component-based clustering. Neurons were recorded without pre-selection, and only trials with stable firing patterns were included in the analysis.

### Working Memory Task

We trained monkeys on three types of working memory tasks: spatial, feature, and conjunction. In all tasks, stimuli were presented within a 3 × 3 grid of positions spaced 10° apart. Spatial stimuli were 2° white squares; only the eight peripheral positions were analyzed. Shape stimuli were 2° geometric shapes (e.g., circle, diamond, H, plus, triangle).

Each trial began with a 1-s fixation period, followed by a 500-ms cue presentation, a 1.5-s delay, a 500-ms test-stimulus presentation, and a second 1.5-s delay. In spatial tasks, the test stimulus appeared either in the same or opposite location to the cue. In feature tasks, the test stimulus always appeared at the same location and either matched or differed from the cue in shape. After the second delay period, monkeys indicated whether the cue and test stimulus matched by making an eye movement (saccade) toward one of two colored targets (green= match, blue= nonmatch), whose positions were randomized to prevent location-based guessing.

In the conjunction task, both shape and location had to match. Only four shape-location combinations were used per session, making this task the most demanding due to the need to maintain two features in memory simultaneously.

### Data Analysis

We did all the analysis and modeling in MATLAB. For each neuron, we computed mean firing rates across task epochs: fixation, cue, first delay, test stimulus, and second delay. We defined neurons as task-responsive if their firing rate in any epoch exceeded the fixation firing rate (one-tailed paired t test, α = 0.05). We defined persistent neurons as neurons with a suprathreshold firing during the first delay that exceeded the baseline firing rate (35–37).

For each working memory dataset, we recorded firing rates across 8 cues × 12 trials for the spatial and feature tasks, and 4 cues × 24 trials for the conjunction task. This resulted in a total of 96 trials, with *t*_trial_ time points in each trial. We normalized the firing rates of each neuron separately within each trial to prevent any information leakage between the training and test sets during decoding analysis.

To prepare the data for modeling, we concatenated these firing rates across all trials, resulting in a time-series matrix of size *n* × *t*_all_, where *n* is the number of neurons and *t*_all_ is the total number of time points across all trials. It follows that the length of the full time-series was *t*_all_ = 96 × *t*_trial_ time points.

To track the experimental conditions throughout the time series, we generated two sets of labels for downstream analysis and modeling. The first set labeled each time point with the trial identity. The second set labeled each time point with one of the seven task epochs: pre-fixation, fixation, cue, first delay, test stimulus, second delay, or response.

### Decoding Analysis

We decoded the stimulus identity (location or shape) using an SVM classifier. We used a sliding window of 100 ms with a step size of 25 ms to extract neuronal firing rates at each time point. For each decoding iteration, we randomly selected 200 neurons from either the persistent or non-persistent neuronal populations and evaluated the performance with 10-fold cross-validation. We repeated this process across the full time-course of all three match/non-match tasks, for both the empirical and model data. Note that the non-persistent population of the conjunction task contains 211 neurons, so subsamples for that condition necessarily show high overlap and thus low sampling variability.

The chance decoding level was 1/(8 possible stimuli) or 12.5%, for the spatial and feature tasks, and 1/(4 possible stimuli), or 25%, for the conjunction task.

### Between- and within-cue distances of model statistics

We computed model statistics on normalized empirical activity at the delay period (0.75 to 1.75 seconds after cue onset), separately for persistent and non-persistent neurons. For the temporal components, we computed the first three temporal principal components and used Procrustes rotation to align these to a reference principal component analysis of all pooled neurons across trials.

For each statistic, we normalized every feature across the 96 trials, as in the decoding analysis, computed the squared Euclidean distance between all 4,560 pairs of trials, and finally normalized these distances to zero mean and unit standard deviation.

### General Formulation of Nullspace Sampling

We recently developed a nullspace sampling method that can generate synthetic population activity data that satisfy diverse neurobiological constraints. Here, we briefly describe the key aspects of this method. We refer readers to our previous studies for additional details (22,23).

We use **X** ∈ ℝ^*n*×*t*^ to denote the empirical data, and 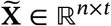 to denote one sample of synthetic data. Correspondingly, we use **x**_*i*_ ∈ ℝ^*n*×1^ and 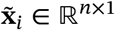 to denote columns of **X** and 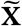 at specific time points. Each column thus denotes the activity of all neurons at each time point.

We used nullspace sampling to generate synthetic samples 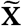 by sequentially generating 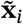 that each satisfies a set of linear constraints, encoded by the matrix **A**_*i*_ ∈ ℝ^*m*×*n*^. Mathematically, we sequentially sampled 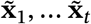 that each satisfied

,
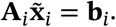

In practice, we can define each sample 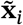 as

,
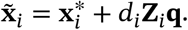

We now describe each of the terms in this equation.

First, 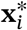 is a unique minimum-norm solution of **A_*i*_ x_*i*_** = **b_*i*_**, given by 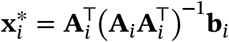.

Second, **Z**_*i*_ ∈ ℝ^*n*×*k*^ is an orthonormal basis of the nullspace of **A**_*i*_ such that **A**_*i*_**Z**_*i*_ = **0**_*m*×*k*_ and **Z**^⊤^**Z**_*i*_ = **I**_*k*_, where **I**_*k*_ is the *k* × *k* identity matrix.

Third, **q** ∈ ℝ^*k*×1^ is a unit-norm random weighting vector, and together with **Z**_*i*_ allows us to generate, uniformly at random, vectors 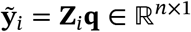 that satisfy 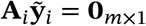. Finally, *d*_*i*_ is a scaling param eter that we use to set our solutions to have target norm.

We built on this basic formalism to implement specific models that satisfy principal components, cor relation structure, and mean firing rate, as we detail below.

### Cluster and component models

We defined the tuned-cluster mean model by first clustering neurons based on cue preference, defined as the cue eliciting the largest mean response during the cue epoch relative to the fixation epoch. We then preserved the mean neuronal activity within each cluster, separately for the persistent and non persistent categories. Specifically, for each neuronal cluster *h* and time point *i*, we sampled cluster activity vectors 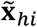that preserved the empirical cluster mean and norm:

,
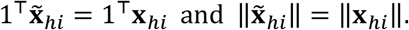

We defined the temporal component models by first dividing the time-series data into two subsets based on persistent and non-persistent neuron types. For each subset, for every single trial (each of the 96 trials), we computed the first three principal components across time to capture the low-dimensional temporal structure of population activity, including persistent activity. We then used nullspace sampling to generate synthetic data preserving the same dominant low-dimensional structure of persistent activity.

To do this, we normalized each column of X to zero mean and unit norm, and worked with the empirical temporal correlation matrix **C** = **X**^⊤^**X** ∈ ℝ^*t*×*t*^ and its eigen-decomposition

,
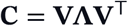

into **V** ∈ ℝ^*t*×*t*^, a matrix containing its eigenvectors as columns, and **Λ** ∈ ℝ^*t*×*t*^, a diagonal matrix containing the corresponding non-negative eigenvalues, ordered by magnitude.

We now define a synthetic correlation matrix,

,
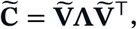

which has identical eigenvalues, but a modified matrix of eigenvectors 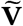.

Let us now consider the structure of 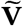. We know that the first *k* columns of this matrix must be identical to the columns of **V**, by the definition of the constraints. It follows that, to generate our synthetic correlation matrix of interest, we must sample the remaining (*k* + 1) to *t* eigenvectors of 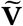.

We can do this by sequentially sampling 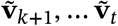, such that each 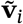 in this sample satisfies orthogonality constraints,

,
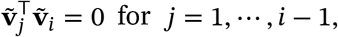

and unit-norm constraints,

,
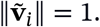

We can do this by translating the problem into the formalism of nullspace sampling. For each vector 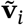, we can first define a system of linear equations,

,
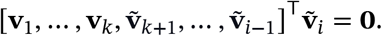

Once we have 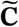, our task reduces to the sampling of time series 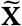 that satisfy

,
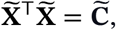

and that therefore have the same leading principal components as **X**. We can do this by sequentially sampling the columns of this matrix, 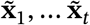. We first sampled 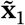 to be a normally distributed random vector with zero mean and unit norm. We then sampled the vectors 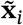 for *i* = 2, …, *t* by defining a system of equations,

,
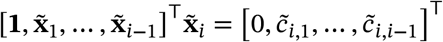

where 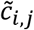 denote the (*i, j)*^th^element of 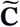, 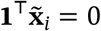 enforces zero mean, and 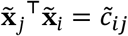 ultimately enforces the correlation coefficient (because all 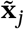 have zero mean and unit norm). Finally, we rescaled and shifted the sampled columns to restore the empirical mean and norm of each trial. See our recent paper for additional details (22).

### Epoch Correlation models

As described in the data analysis section, we labeled neurons as either persistent or non-persistent. Separately, we associated each time point in the data with one of 96 trials and one of seven task epochs (pre-fixation, fixation, cue, first delay, test stimulus, second delay, and response). This process produced a total of *m* = 96 × 7 = 672 unique task epochs. Each epoch therefore represents the activity of all neurons at a cue-specific trial.

In this section, we constructed a model that captures the temporal structure of the working memory task paradigm by constraining the pairwise correlation of these task epochs. We did this separately for persistent and non-persistent neurons to distinguish their respective activation patterns and to isolate each group, minimizing crosstalk and indirect influences between them.

To build this model, we first computed an unnormalized centroid matrix **S** ∈ ℝ^*n*×*m*^. Each column of this matrix encoded the summed activity of neurons within each epoch-specific trial, and the correlations of the persistent and non-persistent subsets of this matrix encoded our temporal constraints of interest.

We generated a synthetic time-series matrix 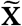 in two steps. In the first step, we generated 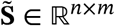 a synthetic matrix of unnormalized centroids that satisfied the correlation constraints separately for the subsets of persistent and non-persistent neurons. We did this by using the same algorithm for sampling data described in the previous section. For the all-epoch correlation model, the constraints imposed on 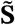 ensured that the complete *m* × *m* correlation structure of 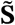 matches the empirical. However, for the same-epoch correlation model, this condition was applied only to the same-epoch correlations between trial pairs (seven matrices, each of size 96 × 96).

We finally converted this unnormalized centroid matrix 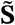 to the full time series matrix by expanding each summed centroid to its constituent time points. We did this by sampling the time points of each epoch *i*, 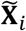, to satisfy 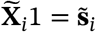.

### Whole-trial correlation model

To serve as a control for correlation models, we constructed a model that preserved coarser features of the data. As above, we split the time-series data into persistent and non-persistent subsets and generated a model constrained separately for each type of neuron.

For each trial time series, we computed the mean neural activity across time, resulting in two matrices of size n_pers_ × 96 and n_non−pers_ × 96. Modeling constraints were then applied to preserve the correlation structure between these 96 trial-averaged neuronal activities, producing two separate 96 × 96 correlation matrices, one for persistent neurons and one for non-persistent neurons.

Unlike the epoch-correlation models, this model does not account for within-trial temporal structure. As a result, it provides a minimal representation of the dynamics in the original data and serves as a control for evaluating the importance of more detailed temporal structure.

