## Supplementary Figures for "Disentanglement of persistent and dynamic neural substrates of working memory"

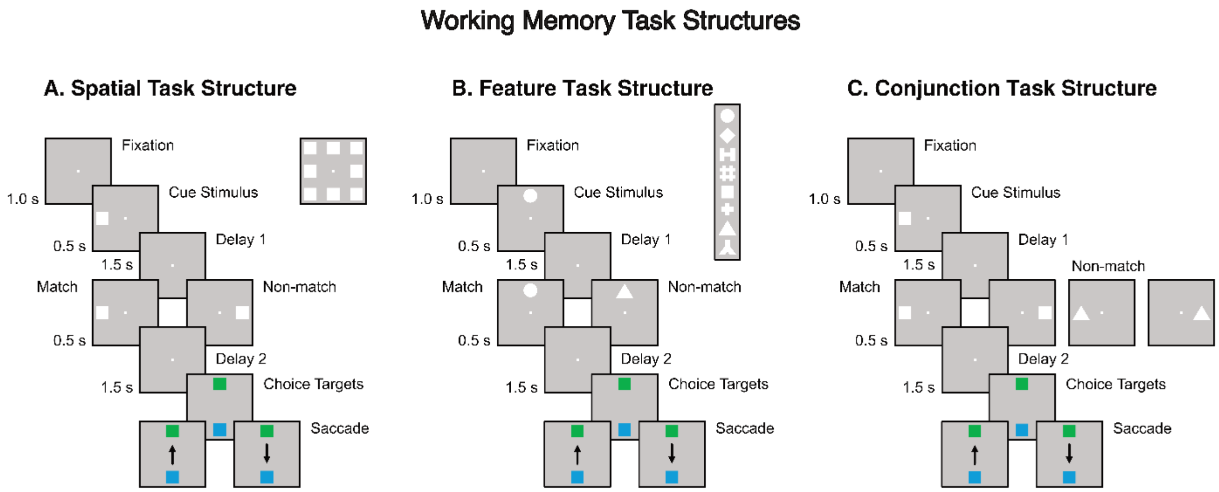


#### Figure S1. Structure of the three match/non-match working-memory tasks.

Monkeys performed three variants of a delayed match-to-sample task while maintaining central fixation throughout each trial. At the end of each trial, the monkeys indicated whether the cue and test stimuli matched by making a saccade: either to a green target if the stimuli matched, or to a blue diametrically opposed target if they did not.

**A.** Spatial task (all stimuli defined by location). The inset shows the eight stimulus locations used for analysis.

**B.** Feature task (all stimuli defined by shape). The inset shows the eight stimulus shapes used for analysis.

**C.** Conjunction task (all stimuli defined by both location and shape, with two locations and two shapes per session).

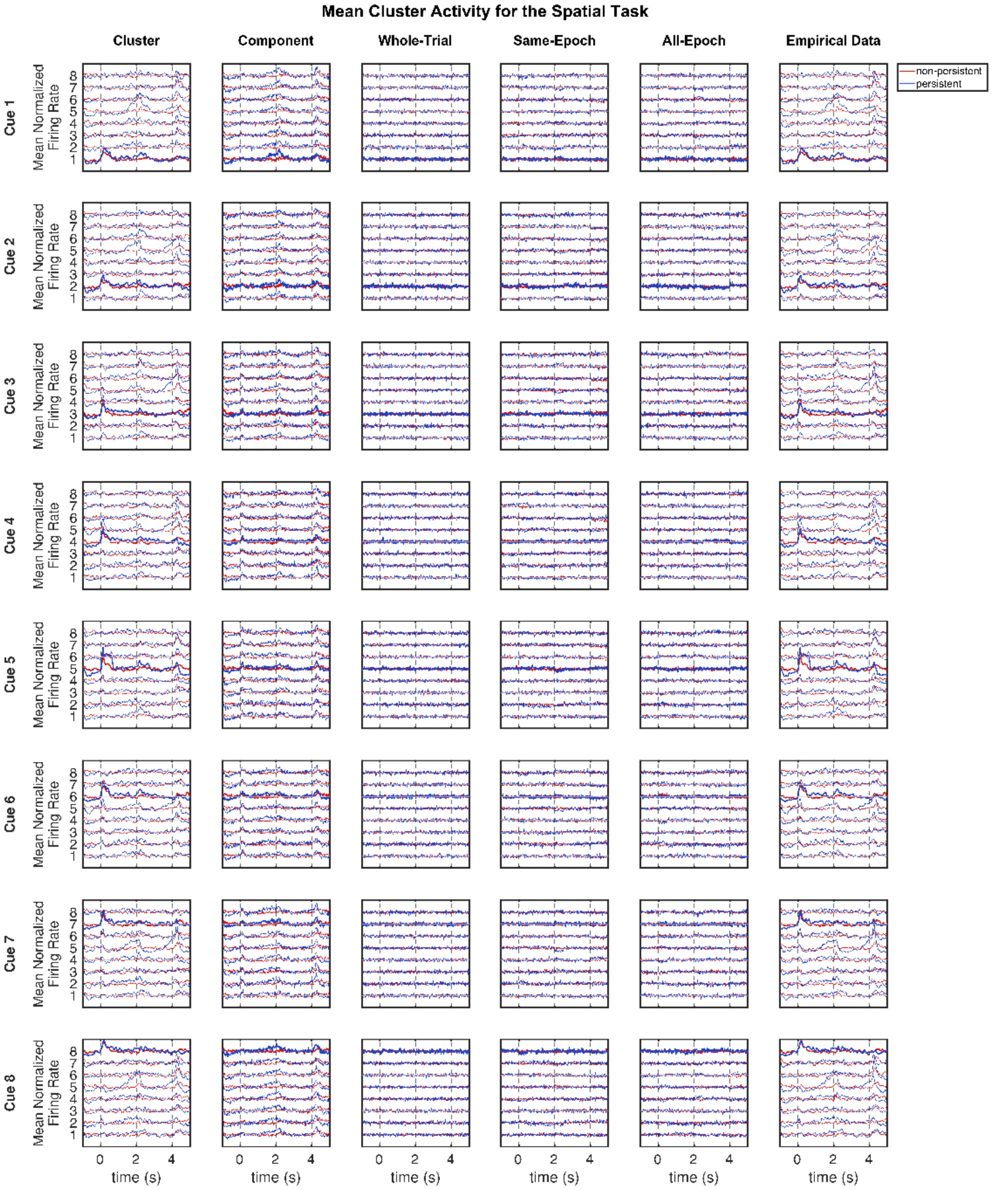


**Figure S2. Average population activity of the preferred-cue neuronal clusters for all cue stimuli in the spatial task.**

Rows show individual cue stimuli. Columns 1–2 show statistics of the cluster and component-model data. Columns 3–5 show statistics of correlation-model data. Column 6 shows statistics of empirical data. Within each panel, traces show the mean normalized firing rate of each preferred-cue neuronal cluster, offset vertically by cluster, with the cluster matching that row's cue drawn in bold. Blue traces show activity of persistent neurons and red traces show activity of non-persistent neurons. Dashed lines at 0, 2, and 4 s denote the onsets of the cue, the test stimulus, and the response epoch, respectively.

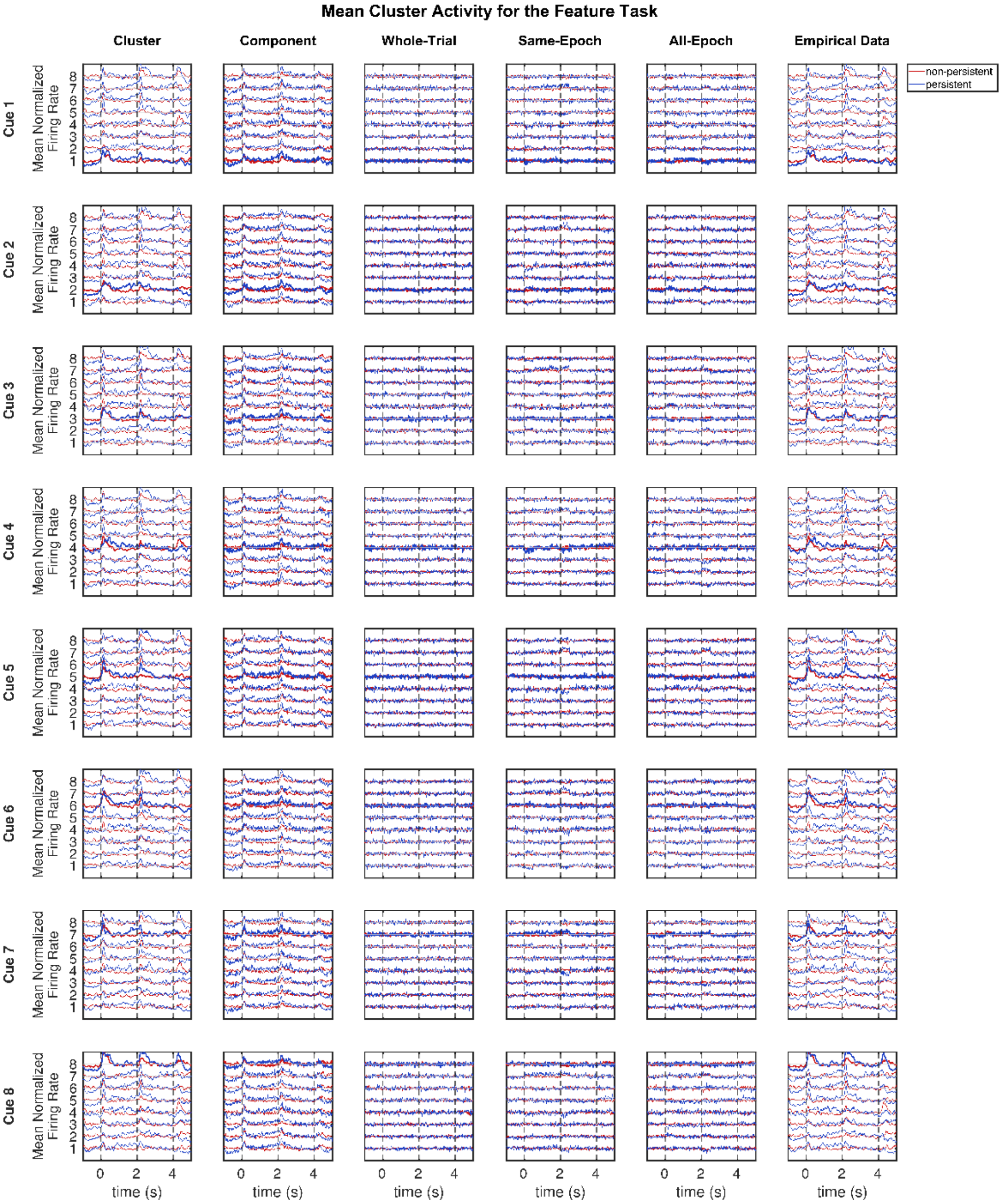


**Figure S3. Average population activity of the preferred-cue neuronal clusters for all cue stimuli in the feature task.**

See Figure S2 caption for details.

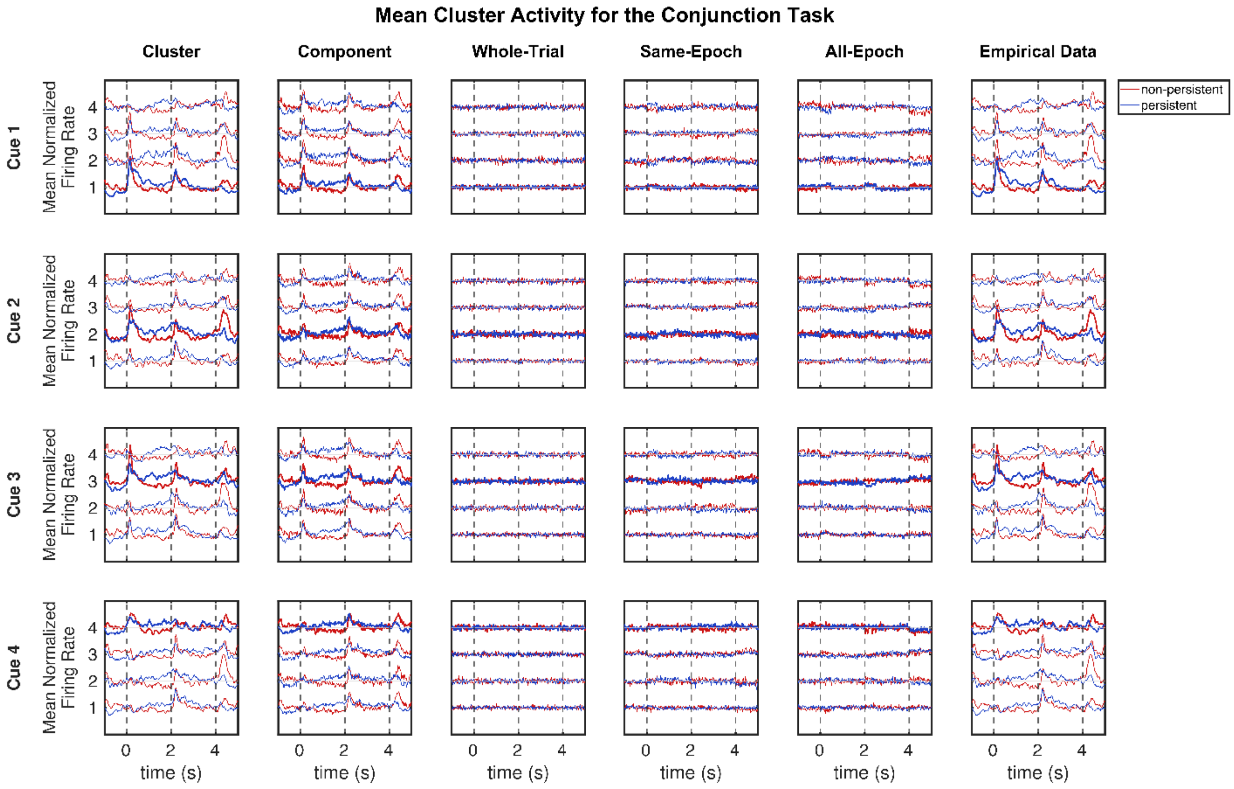


#### Figure S4. Average population activity of the preferred-cue neuronal clusters for all cue stimuli in the conjunction task.

See Figure S2 caption for details.

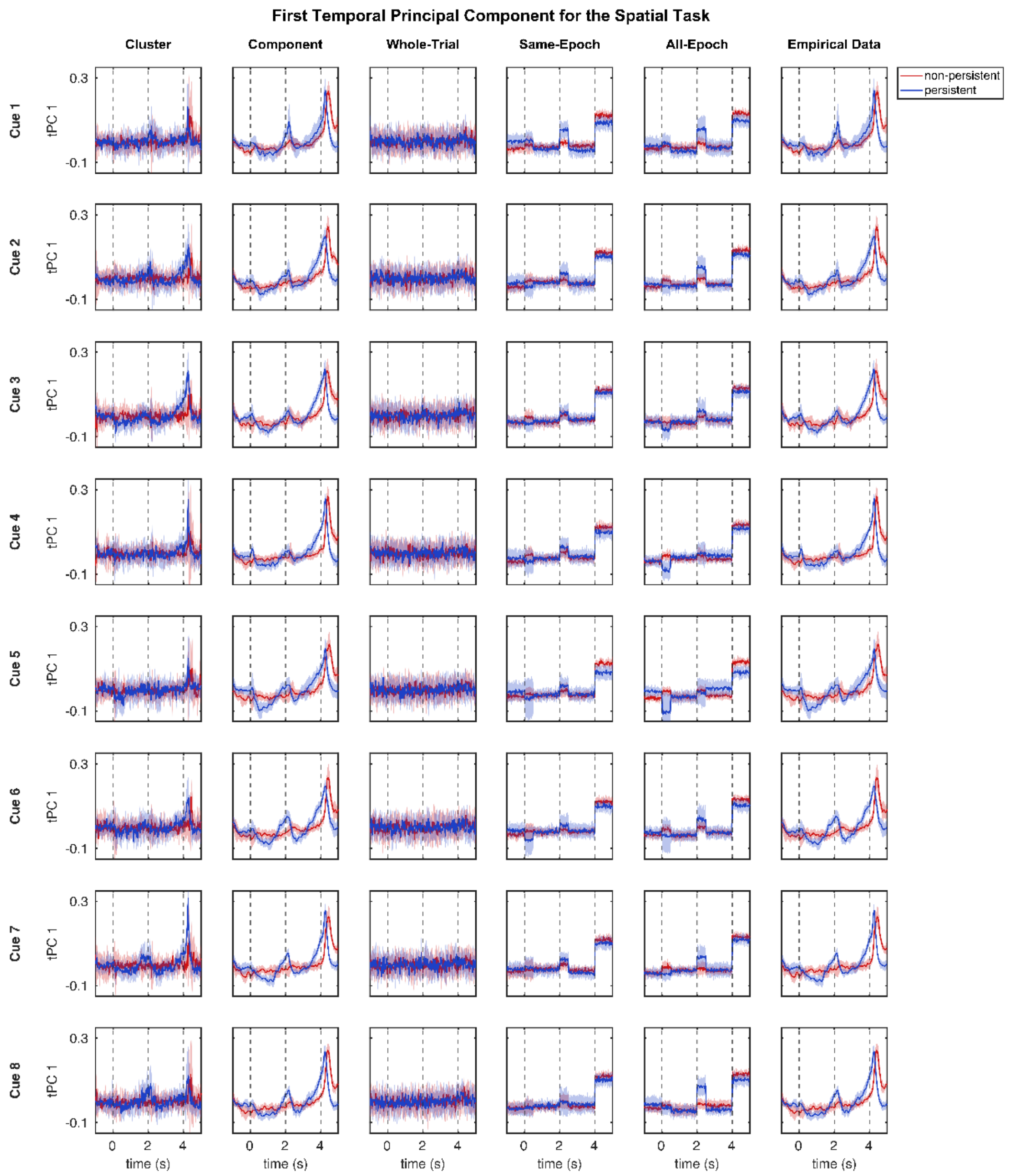


### Figure S5. First temporal principal component for all cue stimuli in the spatial task.

Rows show individual cue stimuli. Columns 1–2 show statistics of cluster and component-model data. Columns 3–5 show statistics of correlation-model data. Column 6 shows statistics of empirical data. The shaded area shows ± 1 standard deviation across the 12 trials. Red denotes non-persistent and blue denotes persistent neurons. Components were aligned to a common reference by orthogonal Procrustes before averaging. Dashed lines at 0, 2, and 4 s denote the onsets of the cue, the test stimulus, and the response epoch, respectively.

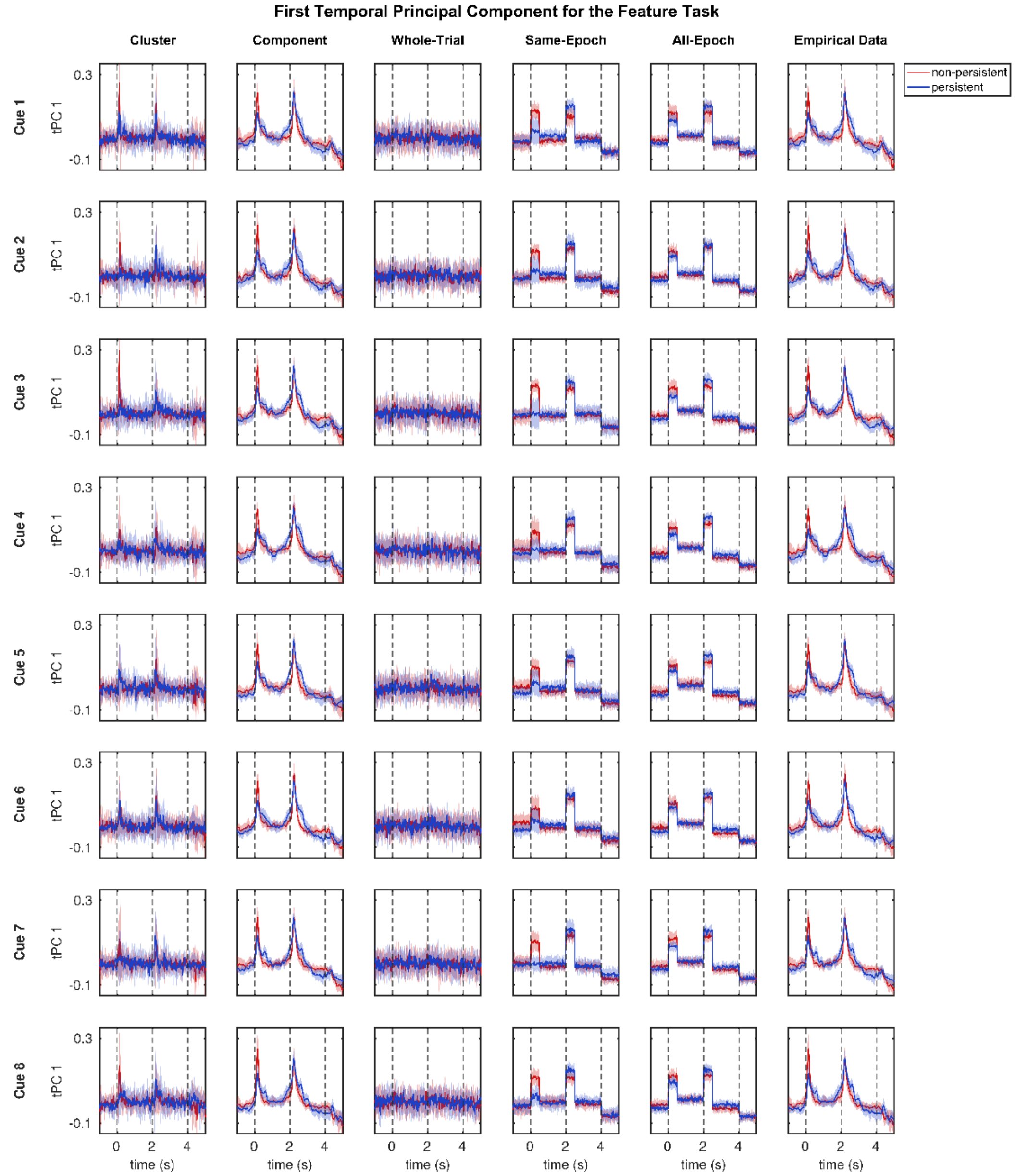


### Figure S6. First temporal principal component for all cue stimuli in the feature task.

See Figure S5 caption for details.

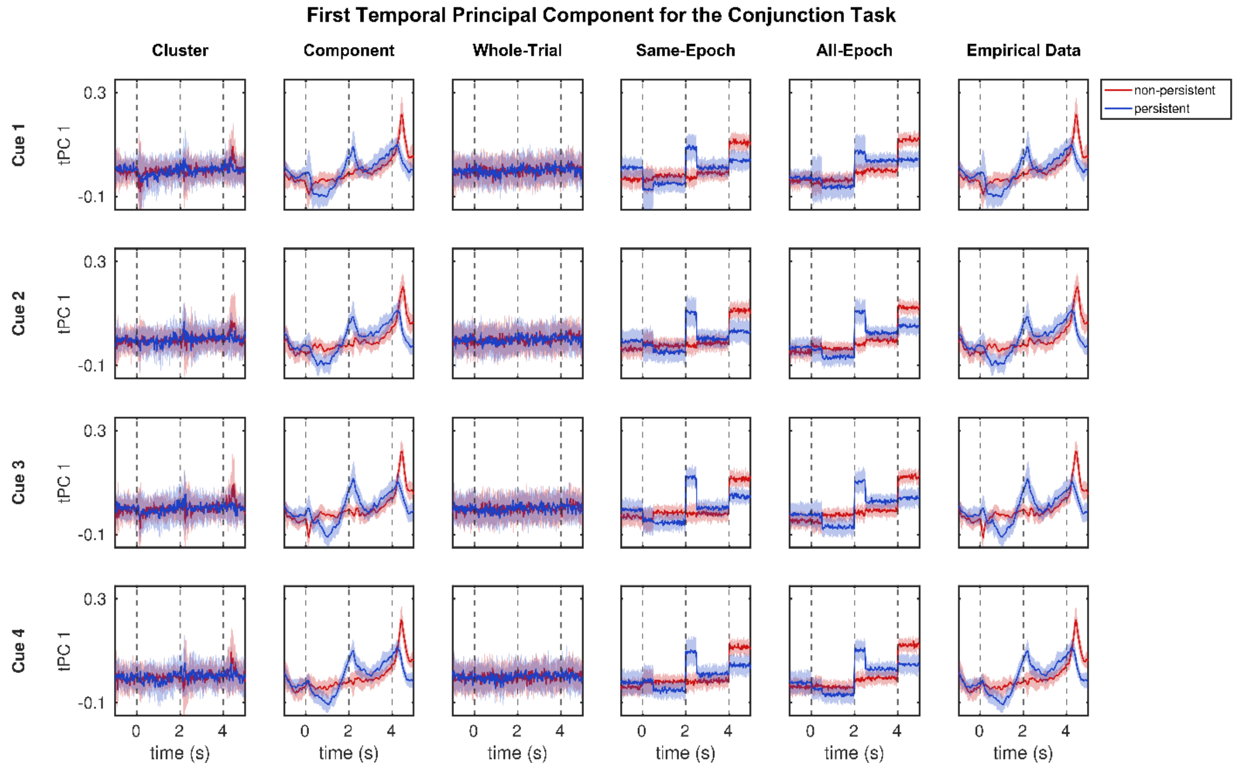


**Figure S7. First temporal principal component for all cue stimuli in the conjunction task.**

See Figure S5 caption for details.

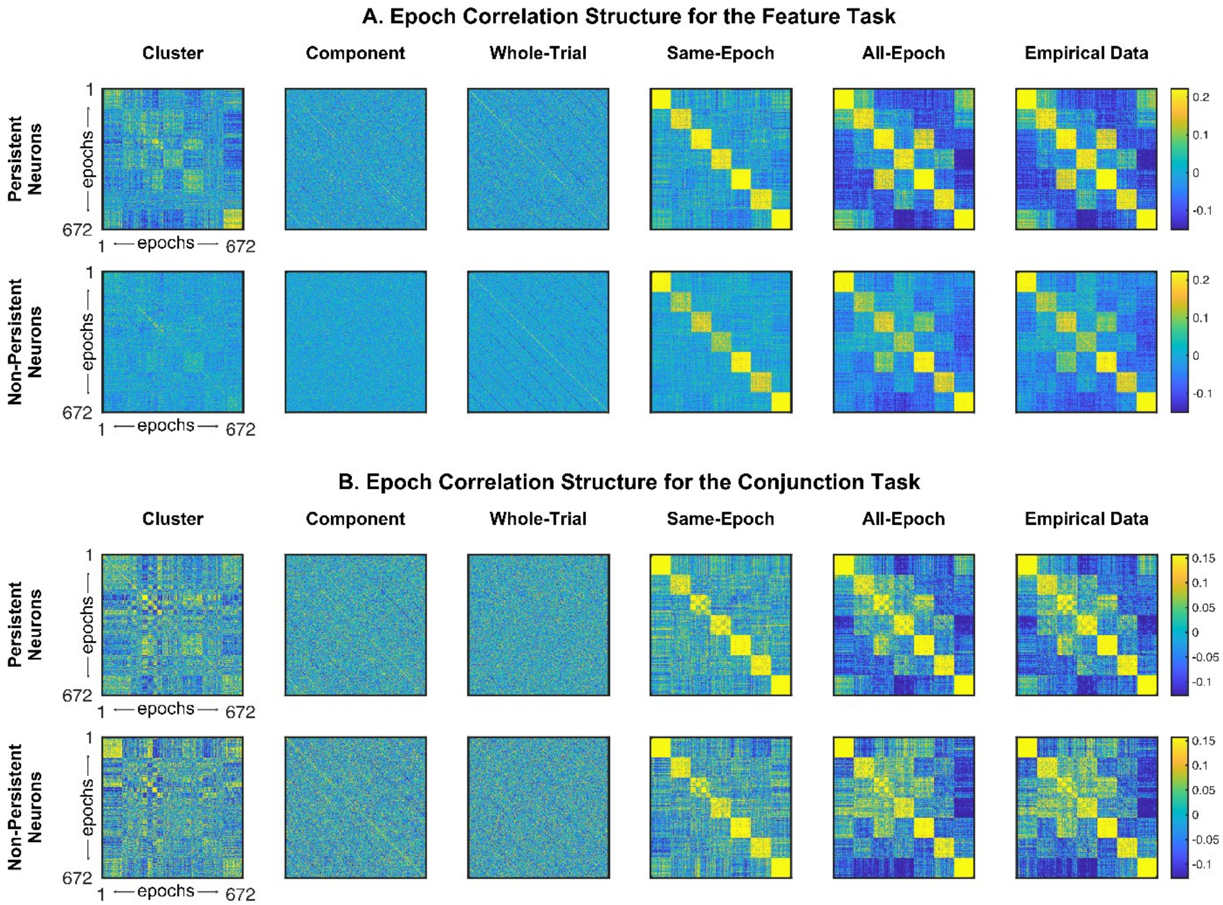


#### Figure S8. Epoch correlation structure for feature and conjunction tasks.

Pairwise correlations between mean activities within each epoch across all trials in the feature (**A**) and conjunction (**B**) tasks, shown separately for persistent and non-persistent neurons and ordered as in **Figure 2C**.

Columns 1–2 show cluster and component-model data. Columns 3–5 show correlation-model data. Column 6 shows empirical data.
